# Expanding the access of wearable silicone wristbands in community-engaged research through best practices in data analysis and integration

**DOI:** 10.1101/2023.09.29.560217

**Authors:** Lisa M. Bramer, Holly M. Dixon, David J. Degnan, Diana Rohlman, Julie B. Herbstman, Kim A. Anderson, Katrina M. Waters

## Abstract

Wearable silicone wristbands are a rapidly growing exposure assessment technology that offer researchers the ability to study previously inaccessible cohorts and have the potential to provide a more comprehensive picture of chemical exposure within diverse communities. However, there are no established best practices for analyzing the data within a study or across multiple studies, thereby limiting impact and access of these data for larger meta-analyses. We utilize data from three studies, from over 600 wristbands worn by participants in New York City and Eugene, Oregon, to present a first-of-its-kind manuscript detailing wristband data properties. We further discuss and provide concrete examples of key areas and considerations in common statistical modeling methods where best practices must be established to enable meta-analyses and integration of data from multiple studies. Finally, we detail important and challenging aspects of machine learning, meta-analysis, and data integration that researchers will face in order to extend beyond the limited scope of individual studies focused on specific populations.

## 1. Introduction

Silicone wearables as passive sampling devices have emerged as a powerful and versatile personalized exposure assessment tool, allowing researchers to characterize chemical exposures for a wide variety of organic chemicals and study the impact of exposures on human health [1]. The use of silicone wearables in research, especially wristbands, has grown substantially since the first publication in 2014 [2]. Thousands of participants from several countries on six continents have worn wristbands [3] and there have been over 60 peer-reviewed papers published to date [1]. Since wristbands are easy-to-wear, do not require in-person consultation or training, and can be transported at ambient temperature in the mail back to the laboratory for analysis, they are a convenient choice for researchers and study participants alike even in challenging scenarios like disasters or pandemics [4-6].

However, despite the growing use of wristbands in research, the majority of individual wristband studies are limited due to small sample size and narrow population focus. In addition, no established best practices for analyzing wristband data across multiple studies exist, thereby limiting impact and access of these data for larger meta-analyses. Dixon et al. is the only study that has taken wristbands from multiple studies and reported trends in chemical exposure patterns across the globe [7]. In this paper, authors took wristband extracts from 14 different communities on three continents and re-ran those extracts on the same analytical method for the presence and absence of 1530 chemicals. Authors identified common chemical mixtures between geographically diverse participants. Dixon et al. also reported that wristbands worn in Texas post-Hurricane Harvey had the highest mean number of chemical detections compared with the other study locations, illustrating that comparing wristband studies from a diverse set of communities and geographical areas can highlight populations with unique chemical exposure profiles and therefore unique health risk profiles.

Re-running wristband extracts from different studies on the same analytical method as done in Dixon et al. [7] is not a sustainable strategy for using wristband data to better understand broad exposure patterns and trends. We need new data analysis strategies to combine wristband data from multiple studies or use meta-analysis procedures, which would increase data access and interoperability. The growing number of individual wristband studies can be leveraged by combining data across studies to uncover patterns about personal chemical exposure, which can lead to new human health discoveries and can be used to direct research, interventions, and policy resources towards communities with higher exposure burdens or unique exposure patterns. In this manuscript, we present key considerations for analyzing wristband data and combining data collected from multiple studies. We use datasets from three studies to highlight challenges associated with data structure and missingness and the consequences of varying analysis techniques and choices between studies, which are often overlooked or not addressed in individual studies.

## 2. Methods

### 2.1 Study design and data collection

Our paper illustrates data analysis and integration challenges using chemical exposure data from 616 wristbands worn by participants in two study cohorts, one in New York City and one in Eugene, Oregon. The New York (NY) wristband data was collected as part of an ongoing longitudinal birth cohort study at the Columbia Center for Children’s Environmental Health. Individuals pregnant with a singleton who were 18 years and older wore a wristband for 48 hours in their third trimester of pregnancy [8]. There are two sets of wristband data from the NY cohort that are included in this report, one set includes 22 wristbands from a pilot study collected between 2013 and 2015 [8] (referred to as “NY Pilot”) and the second set includes 168 wristbands from a larger study between 2015 and 2019 (referred to as “NY”). We also include data from 426 wristbands worn by study participants in Oregon in 2017 and 2018 (referred to as “OR”). Study participants were asked to wear wristbands for seven consecutive days in two seasons (summer and winter), wearing a new wristband each day of the study. Study participants had to be 18 years or older, be diagnosed with mild to moderate asthma, be a current non-smoker, and live near Eugene, Oregon.

All participants provided informed written consent in accordance with the Columbia University Institutional Review Board (IRB; #AAAK6753) for the NY cohort and in accordance with the Oregon State University IRB (#8058) for the OR cohort.

We prepared, cleaned, and extracted all the wristbands as previously described [5]. We also created and analyzed several quality control samples throughout the wristband preparation, transport, and laboratory processing steps, which is described in Dixon et al. [5]. We analyzed the New York wristband extracts for 61 organic chemicals with an Agilent 7890B gas chromatograph (GC) paired with a 7000C triple-quadrupole mass spectrometer (MS/MS) [5]. We analyzed the OR wristband extracts for 94 organic chemicals using an Agilent 7890A GS interfaced with an Agilent 5975B MS. Further analytical details can be found in Anderson et al. [9].

### 2.2 Data Processing

We converted chemical concentrations to moles per gram wristband and applied a log transformation (log_2_ pmol/g wristband). We set the concentration value for a given chemical equal to NA if there was matrix interference (Section 3.1) [5]. We conducted analyses using the statistical software R, version 4.1.2 [10]. For each dataset, we filtered out chemicals which were not detected in any wristbands; this resulted in 53, 44, and 69 chemicals in the NY Pilot, NY, and OR datasets, respectively. We masked chemical names in the results as part of our deidentification process.

## 3. Data Properties

### 3.1 Types of missing and censored data

Missing values in data can arise for a variety of reasons and are handled differently depending on the type of missing data. Missing data types are commonly grouped into three categories: missing completely at random (MCAR), missing at random (MAR), and not missing at random (MNAR). When data are MCAR, the probability of an observation being missing is unrelated to any other observed or unobserved factors. Missing values that can be completely explained by another observed variable or variables are MAR. When data are MNAR, the probability of an observation going missing is related to an unobserved variable or variables. There are two primary types of missing data in wristband studies: observations that are below the limit of detection (LOD) and observations that are impacted by matrix interference (MI). Data values below LOD arise from a combination of all three missing data categories. The absence of a quantifiable peak for a sample and chemical of interest (MAR) will result in a missing annotation and results in a majority of the below LOD missing values. Much less frequently, a human error in data processing, such as deletion of a peak in quantification software (MCAR), or a participant’s failure to comply with study protocols, such as removing a wristband for part of a day resulting in low levels of measured chemicals not representative of true exposure may cause data to be annotated as missing and below LOD. Alternatively, MI occurs when a deuterated surrogate peak, used for quantification, is masked; this can arise when a wristband sample contains compounds from personal care products or sweat (MNAR).

### 3.2 Handling missing data

A majority of statistical and machine learning methods require complete observations (i.e. no missing values). Therefore, to leverage these techniques effectively, researchers must decide how to handle missing data, especially when using chemical concentrations from wristbands in a multivariate manner. One solution is to filter any samples or chemicals that contain missing values. We calculated the percentage of chemicals with complete observations across all samples for each of the three studies. Then, for each study we iteratively removed the sample with the most missing values and recalculated the percentage of chemicals with complete data. We summarized the percentage of chemicals with complete data at varying numbers of wristbands. The percentage of chemicals completely observed across all wristbands drops to 50% with only 4, 2, and 3 wristbands for NY Pilot, NY, and OR, respectively. A total of 13, 16, and 21 wristbands with the fewest missing values result in 25% of chemicals with complete observations for NY Pilot, NY, and OR, respectively. When study sizes grow to more than 165 wristbands for NY and OR, only one chemical is observed across all wristbands.

Further, in targeted analytical methods, there is high confidence in the LODs and information about what chemicals are below LOD in wristband extracts contains meaningful data about what people are not being exposed to or are exposed to in very small amounts [9]. Thus, the large number of missing values in wristband data means data removal approaches will significantly diminish the size and information in the data and may introduce bias if missing observations are MNAR.

An alternative approach is imputation of missing observations. The most common imputation approach taken in wristband studies is the replacement of the below LOD missing values with a constant value, such as half the LOD (e.g. [8, 11-13]). Unlike many other mass spectrometrybased measurement fields (e.g. proteomics), a vast majority of below LOD values are due to the true absence of a quantifiable peak, thus half the LOD values are reasonably close to the true unobservable values. However, imputation using a constant value likely does not reflect the true values if they could be measured and can significantly affect the covariance structure of the data, resulting in differences in common downstream analyses, such as principal component analyses (e.g. [11, 14]). As an example, we ran principal component analysis via projection pursuit (PPCA) [15] on the NY Pilot data without imputing missing values (Fig. 1A) and ran k-means clustering [16] based on the first two principal component scores, setting k=4 based on the optimal number of clusters as determined by evaluating the silhouette score [17]. Additionally, we ran PPCA on the NY Pilot data where missing values below the LOD were imputed with half the LOD (Fig. 1B) with samples colored by the clusters assigned based on PPCA results without imputation. The percentage of variability explained by each component is considerably different for the two analyses, and although some samples clustered similarly, several samples formed much different clusters when PPCA was run with imputed values. For example, samples 15, 16, and 22 cluster at the top left in Fig. 1B and the same behavior is not observed in Fig. 1A. Further, we examined the loadings of each chemical on the first principal component (PC1) as seen in Fig. 1C. Large differences in loadings were observed with many chemicals having very little influence on scores in the non-imputed PPCA but having large positive loadings in the half LOD PPCA.

**Fig. 1.**
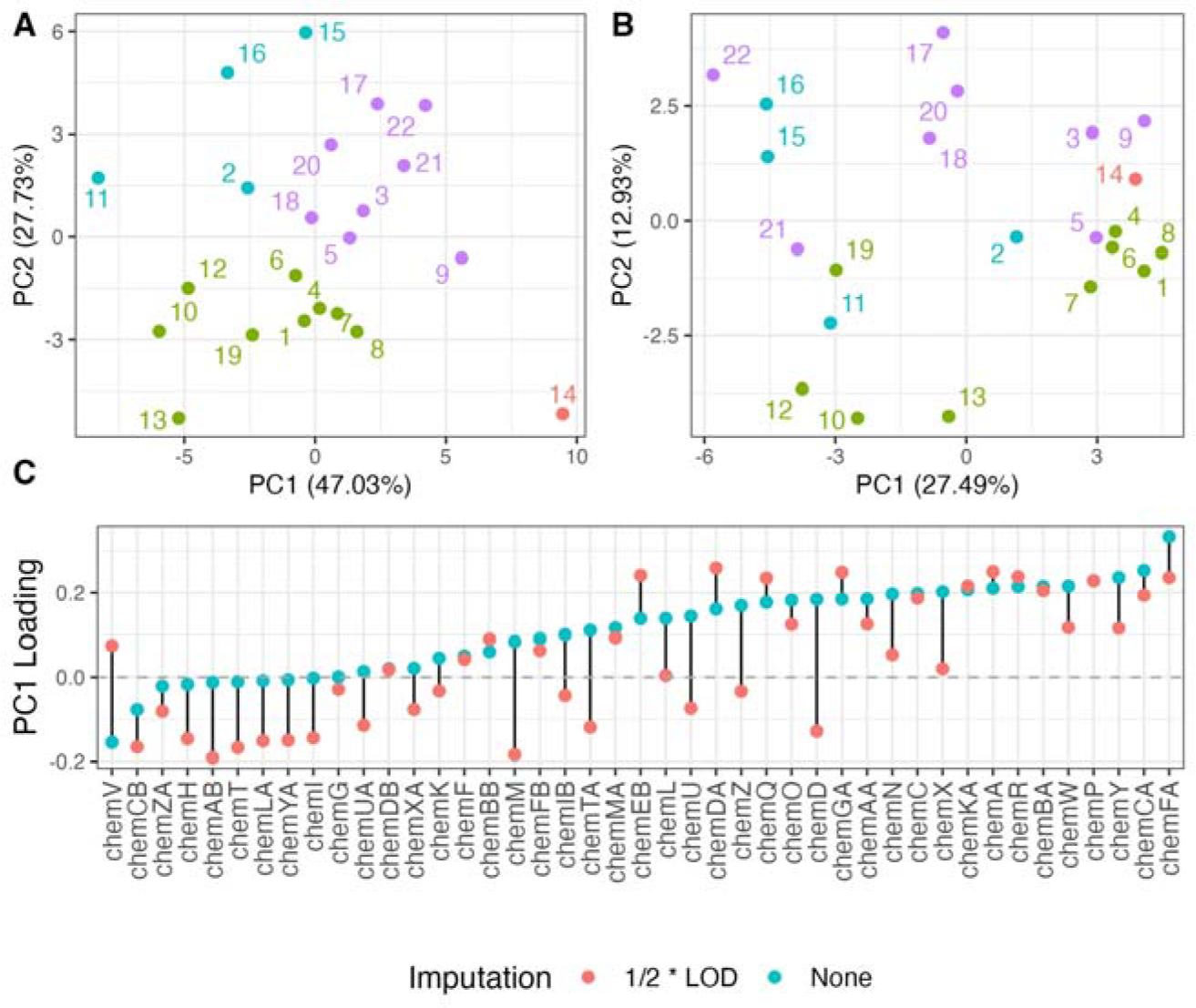
PC1 and PC2 loadings from PPCA for the NY Pilot data for (A) no imputation of missing values and (B) imputation of missing values with half the LOD. Samples are colored by cluster as determined by k-means clustering based on the PC1 scores from PPCA results without imputation. (C) PC1 loadings for each chemical when missing values were not imputed (green dots) and when missing values were imputed with half the LOD (red dots).

Alternatively, missing values can be imputed using more complex algorithms. These methods provide the benefit of introducing variability in imputed values, unlike imputation of half the LOD. A few wristband studies have utilized these approaches to date (e.g. [18]) for all types of missing values. We applied two such example imputation methods to the OR dataset. We imputed missing values using two different methods, random forest imputation [19] and multiple imputation by chained equations (MICE) using the predictive mean metric [20], for chemicals with no more than 40% missing observations. Fig. 2 shows a comparison of imputed values generated by the two methods for observations that were missing due to MI and being below LOD. While some imputed values are very close to one another for the two methods, there are many values that differ by orders of magnitude. Further, the imputation methods were unable to differentiate between the missing value mechanisms, as below LOD and MI observations overlap, nor were they able to impute values below the LOD in nearly all cases. In general, empirically-driven imputation methods are insufficient for imputation of observations below LOD as imputed values are often much larger than the LOD (Fig. 2), which is not consistent with the results of the chemical analysis. Even methods aimed at imputation of left-censored data (e.g. [21]) rely on minimum observed values in the dataset and are still orders of magnitude larger than known LODs, as these algorithms have been designed for different application areas. On the other hand, these imputation methods use the structure of the data to fill in missing observations, and can be useful for resolving missing observations due to MI. The choice of imputation method and underlying assumptions should be carefully considered, as they can lead to significantly different interpretation of a dataset in downstream analyses. Further, no guidance exists nor have any thorough reviews been conducted to determine the threshold of detection rate for a chemical to be included in an analysis and imputed. Researchers have used a wide range of thresholds ranging from 20% [22] to 75% [23] of observations detected for a chemical to be included in downstream analyses. At a minimum, researchers should conduct sensitivity analyses to evaluate the effect of chosen threshold on their results.

**Fig. 2.**
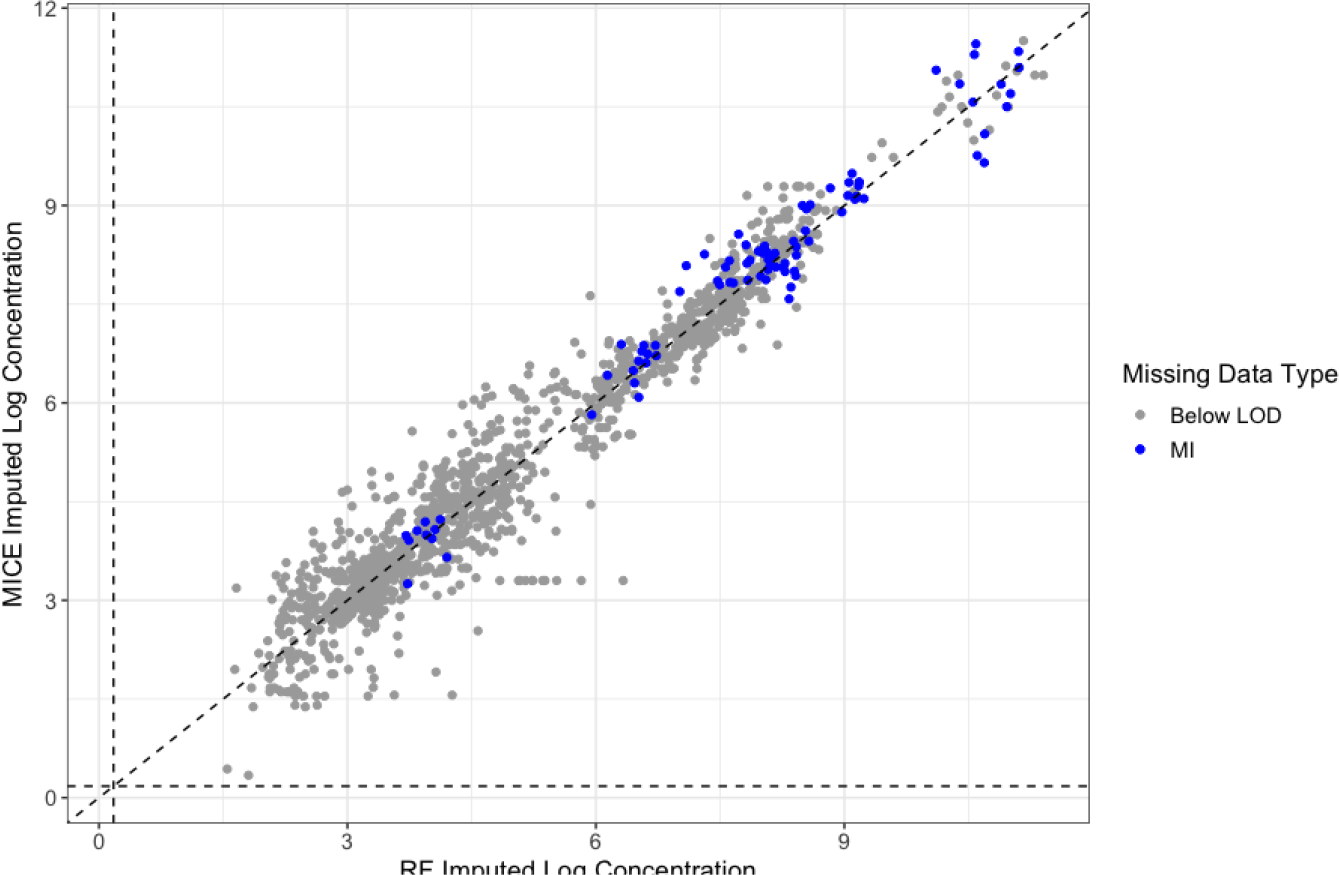
Comparison of MICE and RF imputation methods for two types of missing chemical data, data that is below the LOD (gray dots) and MI (blue dots), in the OR study. The dotted black vertical and horizontal lines represent the median LOD across chemicals.

The final potential solution is to develop or use novel analysis techniques which are tolerant to missing or that can partition sources of variability, so they are not skewed by large numbers of constant values near LOD. For example, in the proteomics field, a statistical analysis technique which combined quantitative and qualitative (presence/absence) models was developed to accommodate and utilize missing observation information [24]. Further research and development of new data analysis models and methods for wristband data is needed to specifically address the challenges described here for wearable wristband data.

### 3.3 Distributional properties of concentrations

Understanding the underlying distribution of chemical concentrations measured by wristbands is of fundamental importance to select appropriate statistical models and analyses to conduct. Fig. 3 shows the distribution of three chemicals from the OR study, with half the LOD filled in for observations below LOD. Within and across chemical observations from wristbands vary by orders of magnitude resulting in distributions heavily skewed to the right (Fig. 3A). Log transformation is a common technique to stabilize variances and transform skewed data distributions to approximately normal distributions and is commonly used in wristband study analyses (e.g. [13, 25]). On a log-transformed scale, chemical concentrations above LOD can be reasonably approximated by a normal distribution (Fig. 3B-D). However, the full distribution of log concentrations is bimodal even for small numbers of observations below LOD, and the distance between observations above and below LOD depends on the chemical.

**Fig. 3.**
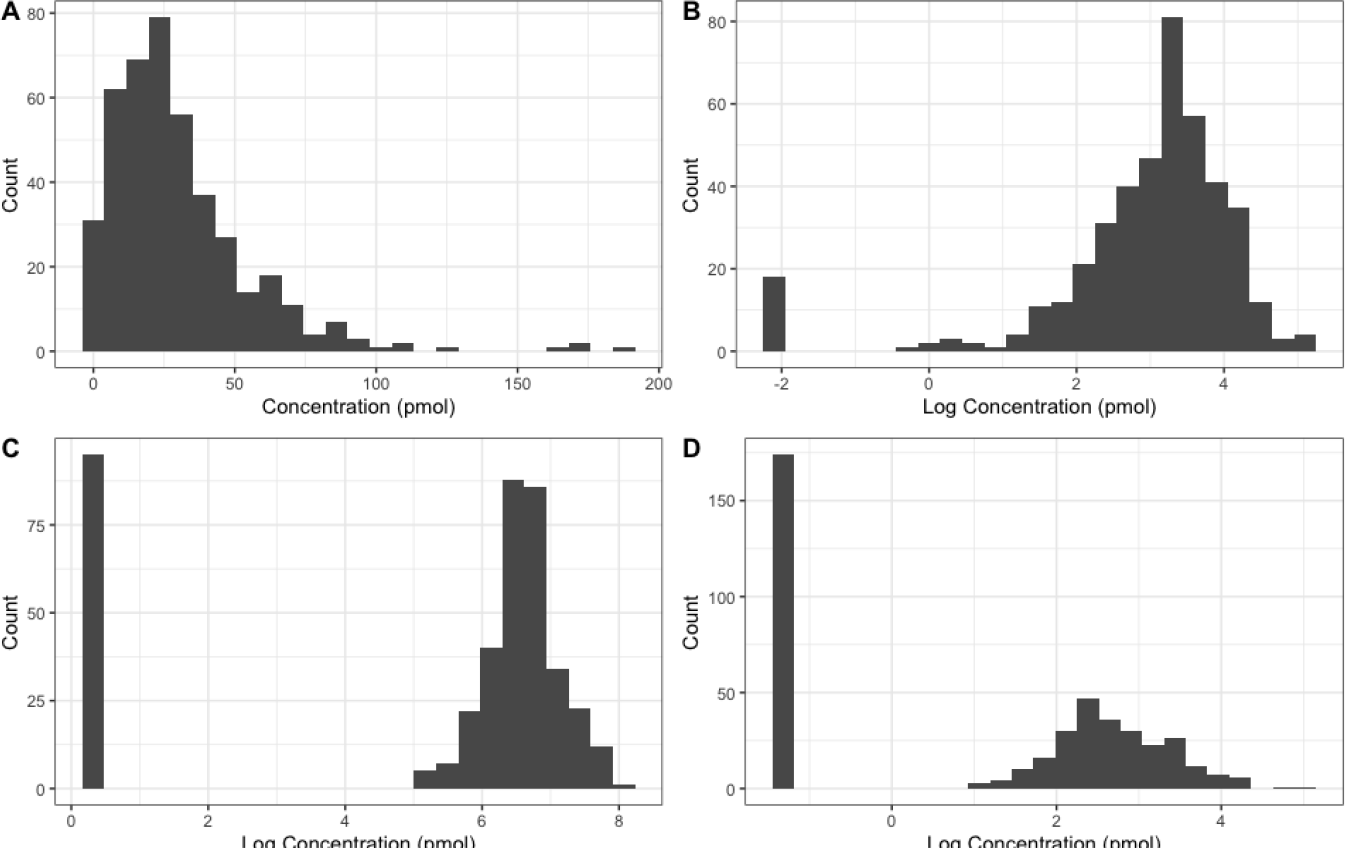
Distribution of three chemicals from the OR study, with half the LOD filled in for observations below LOD (A and B) chemH, (C) chemPB, and (D) chemCB, with chemH visualized on both the (A) linear scale and (B) log scale.

## 4. Data Analysis Methods

### 4.1 Statistical methods and summaries

After conducting a survey of existing literature detailing wristband studies, with a broad range of research applications and hypotheses, and their data analysis methods, several prevalent statistical methods and models emerge, including correlation, linear regression, basic hypothesis testing between groups, and logistic regression. However, despite a small number of statistical models and methods being used in the field, nuances and details of how data preprocessing and model fitting are carried out vary widely. This will create significant problems as the number of wristband studies continues to grow and the research community attempts to combine results from multiple studies through techniques such as meta-analysis [26]. Meta-analysis can be a powerful tool for assessing the consistency and generalizability of results across multiple studies and study populations. However, study combination approaches require consistent statistical data processing and testing procedures.

Perhaps the most common summary statistic used in wristband studies is the calculation of correlations between chemical concentrations measured by wristbands, between a chemical’s concentration profile from wristbands and the profile of another exposure assessment methodology (e.g. urine biomarkers) or to health outcomes. A majority of researchers utilize a non-parametric Spearman correlation, recognizing an assumption of both variables following a normal distribution, as required by metrics such as Pearson correlation, was not appropriate. However, some researchers have calculated Spearman correlation by imputing below LOD observations with half the LOD, or a similar small constant value (e.g. [8, 23, 27, 28]), while other researchers have chosen to calculate correlation using only observations above LOD (e.g. [29]). In the case of linear regression, chemical concentrations are used as the dependent variable or independent variable(s) depending on the research question. Some researchers choose to logtransform concentrations, impute half LOD values for below LOD observations, and limit chemicals used in analyses based on percentage of observations above LOD in an effort to avoid violation of the normality assumption of errors (e.g. [12, 23]). Other researchers restrict linear regression models to chemical concentrations above LOD [30]. Further, the choice of the minimum percentage of detections required per chemical for inclusion in analyses varies widely across studies. When testing for differences in concentrations between groups of interest, some researchers leverage non-parametric tests such as the Wilcoxon Test (e.g. [29]) and others conduct parametric statistics such as a t-test on log-transformed concentrations (e.g. [31]).

Although differences in analyses mentioned above appear small or trivial in nature, the implications on results and conclusions drawn across studies using techniques such as meta-analysis are potentially large. For example, when testing differences in chemical concentrations between groups of interest, a t-test is testing for differences in the mean concentration levels, while the Wilcoxon Test conducts a test for differences in the distribution of values and is typically sensitive to differences in the median, depending on the sample size, but often not the mean unless the sample size is very large [32]. As an illustrative example of the downstream effect of differences in data treatment and modeling, we used the OR dataset. We considered two pairs of chemicals chemUB & chemT and chemH & chemT, for which nearly all wristbands had pairs of observations above LOD (94.8% and 93.7% of 426 wristbands, respectively). We first calculated the Spearman correlation between each pair of chemicals’ concentrations for all complete pairs of observations chemUB & chemT and chemH & chemT as 0.157 and 0.959, respectively. Then for each pair of chemicals, we synthetically introduced missing below LOD values into the data. At each iteration an additional observation was set to below LOD for 1 to 420 (0.2% to 99% of the data) wristbands and the Spearman correlation was calculated 1) with half the LOD imputed for missing values and 2) ignoring missing below LOD observations. Fig. 4 shows the correlation values for the two pairs of chemicals. The treatment of observations below LOD causes the Spearman correlation to differ considerably between the two methods even with small percentages of missing values. As the percentage of detections decreases, correlation calculated using imputation with half LOD inflates and effectively becomes a metric of correspondence between detections and non-detections between the two chemicals rather than measuring the strength of quantitative association, even above thresholds of filtering seen in literature (e.g. 70% marked by the vertical dashed line in Fig. 4). When ignoring missing values from the Spearman correlation calculation, values are centered around the true correlation value but have high variability as the percentage of below LOD observations gets larger. Fig. 4 clearly shows that these two methods for computing correlation are representative of different properties of the data when not all observations are above LOD.

**Fig. 4.**
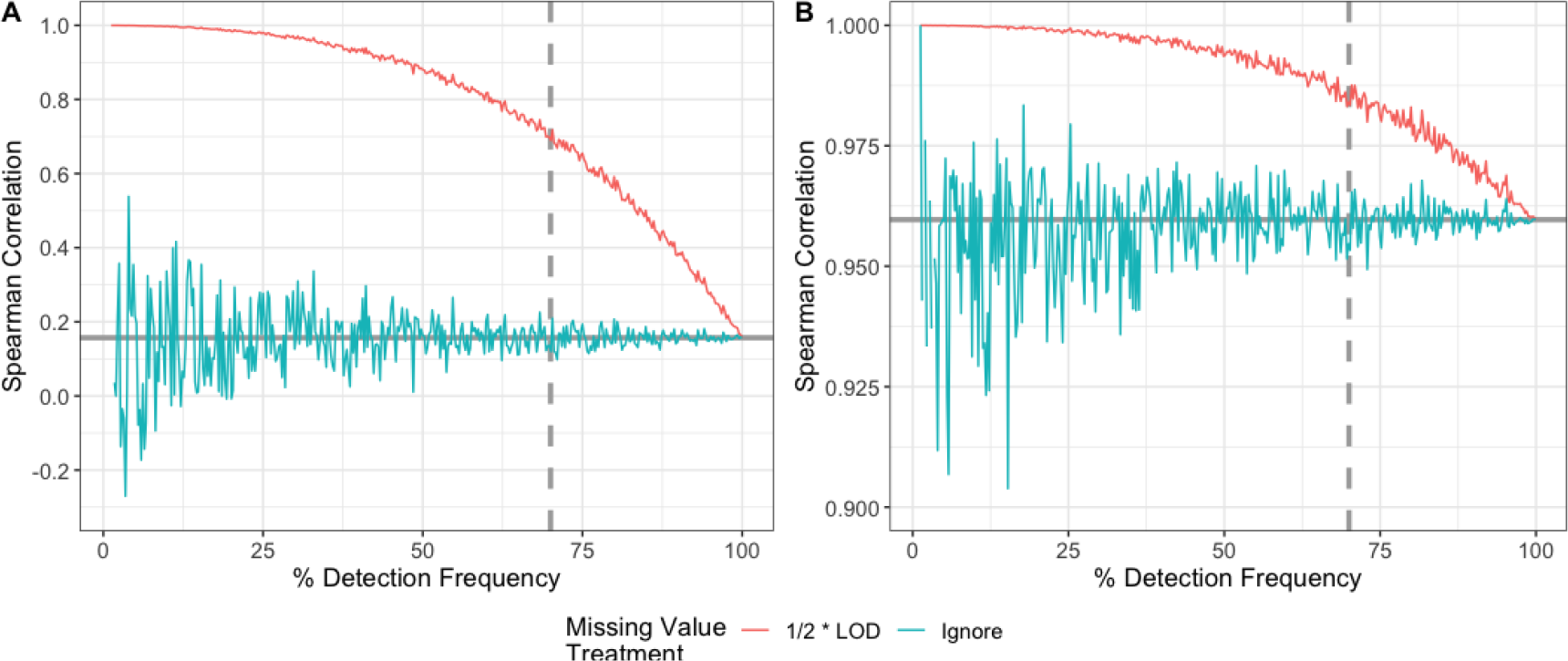
Spearman correlation for (A) chemUB & chemT and (B) chemH & chemT at varying levels of simulated missing values. Vertical lines indicate a common threshold for filtering at 70% detection, and horizontal lines represent the correlation with complete data.

### 4.2 Utilizing machine learning

Machine learning (ML) offers a promising avenue for advanced multivariate analyses, such as discovery of associations between multiple chemicals and a particular health outcome or prediction of a chemical exposure level based on behavior and environmental factors. Despite the promise of ML models, they are most powerful in cases where large sample sizes are available. As the technology grows in popularity and laboratories become more established in the methodology, many studies such as the OR and NY datasets presented here and other studies (e.g. [33]) are reaching sample sizes where ML is a viable option. One limitation of ML is that a majority of methods require no missing values, requiring researchers to again consider and establish best practices when dealing with missing observations due to being below LOD, MI, etc., for wristband data.

If a researcher uses chemical detection status (i.e. detected or below LOD) as the response variable in their model, it is important to note that ML classification model performance can suffer when outcome category frequencies are highly imbalanced [34]. In this case, ML models learn characteristics of the majority class only when prediction accuracy is being optimized. The use of alternative metrics and techniques such as down sampling and upweighting [35] may help alleviate this issue if large enough sample sizes are available. When using a quantitative outcome, a large majority of ML methods, particularly those used on smaller sample sizes (e.g. <1,000), assume that the response variable approximately follows a normal distribution. Therefore, if a researcher uses chemical concentrations as the response variable and imputes below LOD observations, methods such as discriminant analysis, naive Bayes, and support vector machines will be inappropriate. Tree-based methods such as regression trees [36] and random forest regression [37] do not make distributional assumptions and provide more promise for use with wristband data. However, even for these models, the ratio of detects and non-detects, the distance between LOD and detected values, and the optimization or loss function used must be considered carefully. For example, if the mean-squared error is used the model can effectively become a classification model between detections and non-detections with no ability to differentiate large differences in concentrations, because they are still much smaller than the large distance between LOD and observed concentrations. If chemical concentrations are used as predictor variables, some ML also assume normally distributed explanatory variables (e.g. discriminant analysis). Random forest regression models utilize resampling with replacement to grow multiple regression trees. The importance of sample size and even distributional properties, such as number of below LOD observations, becomes important as the resampling method may have a difficult time representing the underlying distribution well, particularly for bimodal distributions. Fig. 5 shows the original distribution of concentrations with half LOD imputed values for chemZ from the NY Pilot dataset. Blue densities show 25 resampled distributions drawn by the random forest model. Some random draws represent the original distribution well, while others do not sample any below LOD observations at all, because of the small sample size. Additional research into nuances of ML methods and establishment of best practices is of fundamental importance as study sizes grow and combination of study datasets becomes possible.

**Fig. 5.**
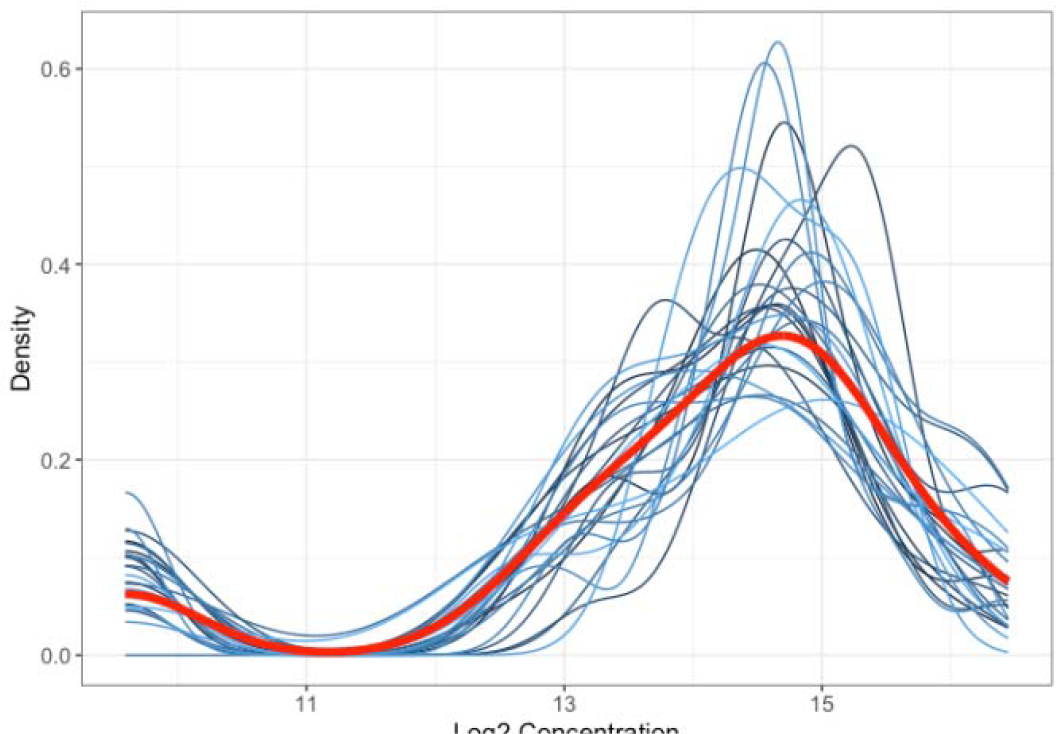
Density of log_2_ chemZ concentrations from NY Pilot data (red) and densities from 25 random forest resample draws (blue).

### 5. Utilizing Data from Multiple Studies

Combining data from multiple wristband studies is crucial to allow researchers to uncover patterns of personal chemical exposure that correlate with potential health impacts across diverse communities. A shared understanding of fundamental wristband data properties and a set of statistically sound strategies to analyze the data is needed in order to pave the way for combining data from multiple studies. However, there are also several other factors that hinder combining wristband data across studies [1, 38, 39]; some are due to research gaps and some are due to differences in how researchers report their data. For example, the way that authors report chemical concentrations varies. Some authors report chemical mass per entire wristband (e.g. ng/wristband) and others report chemical mass per unit mass of the wristband (e.g. ng/g wristband or pmol/g wristband) [1]. When these concentration reporting differences are present and wristband masses are not reported alongside the data, then this inhibits the comparison of chemical concentrations across studies [39]. In addition, more research is needed to understand the variables that influence the rate and amount of chemicals entering into wristbands [1], which could lead to strategies to normalize wristband data across studies where wristbands were worn for different lengths of time and in different environmental conditions. As described in Samon et al., chemical uptake into silicone wristbands is not consistent over time and is dependent on each chemical’s physical-chemical properties and environmental conditions during the study [1]. Silicone post-deployment cleaning and extraction methods also differ between laboratories [3, 39] and it is unknown how these differences may affect quantified concentrations across studies in different laboratories. To reduce additional sources of variability in the data, researchers need to agree on best practices for communicating study protocols to participants to address potential misconceptions up front. For example, participants may think that if they wear the wristband longer than requested, they are helping the research goal and providing more data, when instead they are introducing more variability and complicating interpretation of study results.

While further research into sources of variability due to differences in study designs is needed, how chemical concentrations might be normalized across studies remains an open research question. In the meantime, researchers can consider alternate ways to utilize data from multiple studies. For example, the comparison of study locations can be made by looking for differences in detection frequency, if other important factors are reasonably controlled or equitable between populations. However, different analytical methods for chemical identification and quantification have been developed and used, even for studies coming out of the same research laboratory. In these cases, the chemicals targeted differ. For example, the NY dataset was measured for 61 chemicals, and the OR dataset was measured for 94 chemicals. Of these chemicals, a total of 45 were measured in both studies. The joining of these datasets would result in 110 chemicals in total, and more missing values would be introduced into the joined dataset. However, unlike previous missing data, these missing values would be MAR and downstream statistics would need to account for the additional mechanism for missing values. When comparing the exposure of individuals between study populations, the detection frequency across all chemicals analyzed within each study relative to a measure of central tendency summary across other participants in the same study would give a sense of total exposure level for an individual. Further, many studies have discretized chemical concentration values into categories such as tertiles (e.g. [5, 28]). This concept could be used within a given study to derive chemical tertile profiles for each wristband. Then categorical data analysis strategies such as multiple correspondence analysis [40] could be used to perform clustering of wristband samples across studies to find samples with common patterns. Alternatively, the actual percentile within a study could be recorded and used to visualize and begin to understand exposure patterns across studies. However, the treatment of missing values, either due to a study-based MAR source or a MNAR below LOD source needs to be carefully considered and treated differently. For example, Fig. 6 shows empirical cumulative density curves for chemCB in both the NY and OR datasets with tertile thresholds denoted by gray dashed lines. For this chemical, the proportion of wristbands with values below LOD in the OR study is greater than 0.33. It would be nonsensical to assign some of the wristbands with non-detections to the lower tertile and others to the middle tertile. If all wristbands below LOD were assigned to the lower tertile for OR, a researcher would need to consider if tertiles composed of different proportions of samples are still comparable, and what proportion of LOD observations comparisons are no longer meaningful.

**Fig. 6.**
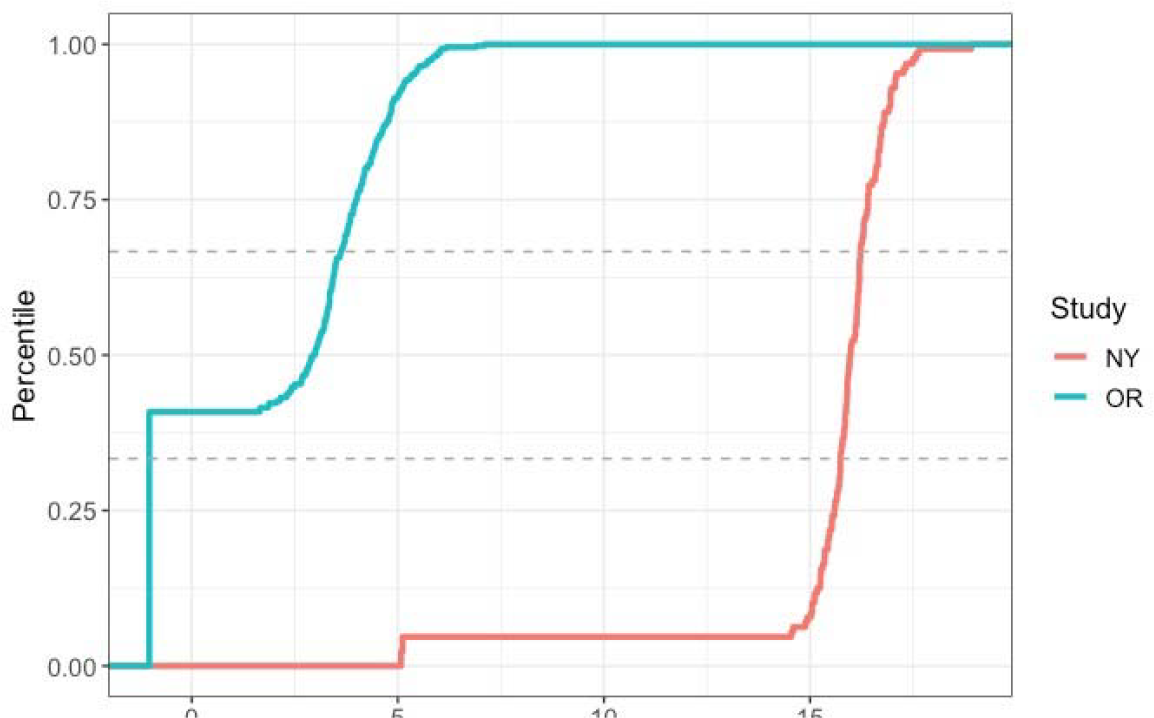
Empirical cumulative density plots of chemCB with tertile thresholds denoted by gray lines.

### 6. Recommendations and Conclusions

The use of silicone wristbands in research studies has rapidly grown over the past few years, especially in community-engaged research [1, 6]. Currently, researchers are using a wide range of data processing and statistical approaches to analyze data from silicone wristbands with a focus on within-study interpretation, but these approaches jeopardize the ability to use the datasets for larger meta-analyses. The need for better guidance and established best practices is evident in the examples shown here, where minor differences in data handling and modeling can lead to vastly different conclusions and interpretation. Some key takeaways and guidance from example analyses presented here are as follows:

- The imputation of half the LOD, or other small constant values, can greatly affect the covariance structure of wristband data (Fig. 1). Even when scaling features, imputation of below LOD values is not recommended as analyses utilizing the covariance structure will be determined by detection rates rather than the intended quantitative information.
- Many dimension reduction techniques for exploratory data analysis have implementations that do not require imputation of missing values, such as projection pursuit PCA. These algorithms are preferable to imputation of below LOD values for wristband data.
- Wristband data chemical concentrations where below LOD values have been imputed with half the LOD do not follow a normal distribution, even when log-transformed, and are bimodal. Methods such as linear regression are not appropriate for this data, and alternative methods such as Gaussian mixture models [41] should be considered.
- Missing value treatment strategies must account for the different missing data mechanisms in wristbands studies, rather than blindly implementing one imputation method on multiple types of missing values (Fig. 2). For example, the use of data-driven imputation algorithms, such as MICE, on below LOD observations can result in imputed values considerably larger than LOD.
- Non-parametric hypothesis tests are not equivalent to parametric hypothesis tests at sample sizes typical of wristband studies. Although methods such as the Wilcoxon Test can be used to skirt the assumption of normally distributed data with half LOD imputed data, even at small percentages of missingness, the tests are effectively determined by detection rates rather than concentration values (Fig. 4).

When presented with data from a wristband study or multiple wristband studies, researchers should first prioritize evaluating the units reported, amount of missing data the effect on the structure of specific chemicals analyzed. If data from multiple studies are present, concentration units should be standardized. Further, only chemicals commonly detected between studies should be considered in downstream analyses. Sensitivity analyses should then be conducted to determine which chemicals should be analyzed using quantitative concentrations or using detected/not-detected information based on the detection frequency of each compound. If data is to be used quantitatively, concentrations should be log-transformed. Tree-based ML methods, which can handle detection information and concentrations simultaneously through LOD imputation, may be considered with large enough sample sizes. Sensitivity analyses looking at the resampled distribution of chemicals should be examined before considering these ML methods. Finally, if the overall exposure profile is of interest, researchers should evaluate if techniques, such as PPCA, provide an interpretable reduced dimension option to represent wristband data. Often in wristband studies, chemicals will all have loadings in the same direction on one of the principal components leading to one potential metric of overall exposure.

This is the first paper to summarize data properties, current data analysis approaches and their issues, and important areas where best practices are needed for wristband data. We demonstrate there is a need for standardized and thorough wristband data analysis methods from the research community, which will create more opportunities to combine wristband data from multiple studies or use meta-analysis procedures, leading to increased data access and interoperability. In addition, more research is needed to understand other factors that hinder the combination of data from individual wristband studies (e.g. how to normalize for differences in wristband wear time and environmental conditions). Overall, a combination of these efforts will enable research to move beyond the narrow population focus of individual studies, leading to new discoveries about personal chemical exposure and potential impacts to human health.

## Acknowledgements

*Preprint of an article published in Pacific Symposium on Biocomputing © 2023 World Scientific Publishing Co*., *Singapore, http://psb.stanford.edu/*.

We thank the study participants for their willingness to engage with our research team. Research reported in this publication was supported by the National Institute of Environmental Health Sciences (NIEHS) under award numbers R21/R33ES024718, P30ES030287, and P42ES016465. The content is solely the responsibility of the authors and does not necessarily represent the official views of the NIEHS. PNNL is a multi-program laboratory operated by Battelle for the U.S. Department of Energy under contract DEAC05-76RL01830.

## Data and Code Availability

- De-identified wristband data is available for download at https://data.pnnl.gov/group/nodes/dataset/33672.
- R code to reproduce all analyses and plots is on GitHub at https://github.com/PNNL-Superfund-Research-Center/PSB_Wristband_Analyses/.

